# Hist2Vec: Kernel-Based Embeddings for Biological Sequence Classification

**DOI:** 10.1101/2023.08.24.554699

**Authors:** Sarwan Ali, Haris Mansoor, Prakash Chourasia, Murray Patterson

**Author notes:** {, }. Equal Contribution (Sarwan Ali, Haris Mansoor, Prakash Chourasia).

## Abstract

Biological sequence classification is vital in various fields, such as genomics and bioinformatics. The advancement and reduced cost of genomic sequencing have brought the attention of researchers for protein and nucleotide sequence classification. Traditional approaches face limitations in capturing the intricate relationships and hierarchical structures inherent in genomic sequences, while numerous machine-learning models have been proposed to tackle this challenge. In this work, we propose Hist2Vec, a novel kernel-based embedding generation approach for capturing sequence similarities. Hist2Vec combines the concept of histogram-based kernel matrices and Gaussian kernel functions. It constructs histogram-based representations using the unique *k*-mers present in the sequences. By leveraging the power of Gaussian kernels, Hist2Vec transforms these representations into high-dimensional feature spaces, preserving important sequence information. Hist2Vec aims to address the limitations of existing methods by capturing sequence similarities in a high-dimensional feature space while providing a robust and efficient framework for classification. We employ kernel Principal Component Analysis (PCA) using standard machine-learning algorithms to generate embedding for efficient classification. Experimental evaluations on protein and nucleotide datasets demonstrate the efficacy of Hist2Vec in achieving high classification accuracy compared to state-of-the-art methods. It outperforms state-of-the-art methods by achieving > 76% and > 83% accuracies for DNA and Protein datasets, respectively. Hist2Vec provides a robust framework for biological sequence classification, enabling better classification and promising avenues for further analysis of biological data.

## 1 Introduction

The rapid advancement of sequencing technology has led to an increase in the quantity of sequence data [16], presenting new opportunities and difficulties for biological sequence analysis. Biological sequence classification, particularly in the domains of protein and nucleotide sequences, is of significant importance in genomics, drug discovery [41], bioinformatics [10,15], and various other research areas [8,26]. Classification of these sequences is crucial in understanding their functions, identifying potential disease-causing variants, and predicting protein structures [5,38,42,11,40]. Traditional classification approaches often rely on feature engineering [1,3,13,23] or sequence alignment methods, having limitations in capturing complex sequence similarities and high-dimensional representations [17,23]. In recent years, kernel methods emerged as powerful techniques for extracting meaningful features and facilitating classification tasks [3]. Motivated by the need for improved biological sequence classification [27,32], this work proposes Hist2Vec, which combines the concept of histogram-based kernel matrices with the effectiveness of Gaussian kernel functions.

Hist2Vec aims to address the limitations of existing methods by capturing sequence similarities in a high-dimensional feature space while providing a robust and efficient framework for classification. By utilizing the unique *k*-mers present in sequences and leveraging the power of kernel methods, Hist2Vec offers a promising avenue for enhancing sequence classification accuracy and enabling further analysis of biological data. We introduce Hist2Vec as a novel kernel-based approach for biological sequence classification. We demonstrate the theoretical properties and advantages of Hist2Vec, including its ability to satisfy Mercer’s condition, ensure continuity, and exhibit the universal approximation property. Additionally, we investigate the practical aspects of Hist2Vec, such as generating kernel matrices, converting these matrices into low-dimensional embeddings using kernel PCA, and applying ML algorithms for sequence classification.

In summary, this paper introduces Hist2Vec as a powerful and effective approach for biological sequence classification. Improved classification accuracy and practical implementation guidelines collectively advance sequence analysis and facilitate a deeper understanding of biological data. Our contributions to this paper are listed below:

1. Introducing Hist2Vec: We propose Hist2Vec, a novel kernel-based approach that combines histogram-based kernel matrices with Gaussian kernels for efficient and accurate biological sequence classification.
2. High-dimensional Feature Representation: Hist2Vec captures sequence similarities by constructing histogram-based representations using the unique *k*-mers present in the sequences. By leveraging the power of Gaussian kernels, Hist2Vec transforms these representations into high-dimensional feature spaces, preserving important sequence information.
3. Improved Classification Accuracy: Through extensive evaluations of protein and nucleotide datasets, we demonstrate that Hist2Vec outperforms state-of-the-art methods in terms of classification accuracy.
4. Advancing Sequence Analysis: Hist2Vec contributes to the advancement of sequence analysis by providing a robust framework for capturing and exploiting sequence similarities. The method offers new insights, fostering further research in genomics, bioinformatics, and related fields.

The remainder of this paper is organized as follows. Section 2 provides a detailed overview of the related work. Section 3 presents the methodology of Hist2Vec, including the computation of histogram-based kernel matrices and the application of Gaussian kernels. Section 4 describes the experimental setup and dataset. Section 5 presents the results of extensively evaluating protein and nucleotide datasets. Finally, Section 6 concludes the paper, summarizes the contributions of Hist2Vec, and discusses future research directions.

## 2 Related Work

Several methods have been proposed for biological sequence classification to capture sequence similarities and facilitate classification [12]. These methods can be broadly categorized into alignment-based methods, feature engineering approaches, and kernel-based methods.

Alignment-based methods, such as BLAST (Basic Local Alignment Search Tool [6]) and Smith-Waterman algorithm [21], rely on sequence alignment techniques to identify similarities between sequences. While these methods effectively detect homologous sequences, they may struggle to capture more complex relationships and handle large-scale datasets efficiently.

Feature engineering approaches involve extracting informative features from sequences and using them for classification [37,22,2]. These features include amino acid composition, dipeptide composition, and physicochemical properties. While these methods are computationally efficient, they often rely on hand-crafted features that may not capture all relevant sequence characteristics.

Kernel-based methods [29,4] can capture complex relationships between sequences. Methods such as the spectrum kernel and string kernel [24] have been proposed for sequence classification, leveraging the concept of *k*-mer frequencies to construct similarity measures. However, these methods may suffer from high computational complexity and memory requirements, limiting their scalability [34]. Kernel methods have been successfully applied to various bioinformatics tasks, including protein fold recognition [31], protein-protein interaction prediction [39], and protein function prediction [7]. These methods leverage the power of kernel functions to map sequences into high-dimensional feature spaces, where the relationships between sequences can be effectively captured.

Mercer’s Condition is a fundamental property of kernel methods, ensuring the kernel matrix is positive semidefinite [28,33]. Many kernel functions, such as the Gaussian kernel, satisfy Mercer’s Condition [35] and have been widely used in bioinformatics applications [25,19]. Hist2Vec addresses the computational complexity and memory requirements by utilizing histogram-based representations of sequences. By counting the frequencies of *k*-mers and constructing histograms, Hist2Vec reduces the dimensionality of the kernel matrix, making it more tractable for large-scale datasets. Furthermore, using the Gaussian kernel in Hist2Vec allows for preserving important sequence information in a high-dimensional feature space.

## 3 Proposed Approach

In this section, we present the proposed approach Hist2Vec. It consists of two main steps: computing histogram-based representations and generating embeddings using kernel PCA.

### 3.1 Histogram-based Representations

A histogram serves as an approximate visual depiction of the distribution pattern of numerical data. Histogram creation involves partitioning the range of values into discrete intervals, commonly called bins, and subsequently tallying the frequency of data points falling within each interval. These bins are defined as consecutive intervals that do not overlap, typically possessing uniform size. By ensuring that adjacent bins are contiguous, histograms effectively eliminate gaps between the rectangular bars, resulting in mutual contact.

Histograms provide an approximate indication of the concentration or density of the underlying data distribution, thereby facilitating the estimation of the probability density function associated with the respective variable. Visually representing the data distribution, histograms depict the frequency or count of observations falling within individual bins. This analytical tool proves valuable in discerning patterns and trends within the data and facilitating comparisons between disparate datasets.

Suppose a dataset *D* = *{x*_1_, *x*_2_, …, *x*_*n*_*} ∈* ℝ^*d*^. The first step of Hist2Vec involves computing histogram-based representations of the input sequences. This process captures the frequencies of specific *k*-mers within the sequences, providing a compact representation that preserves important sequence characteristics.

Given a protein or nucleotide sequence, we compute the *k*-mer spectrum by extracting all possible substrings of length *k*. Each unique *k*-mer represents a distinct feature. We then construct a histogram, where each bin corresponds to a specific *k*-mer and captures its frequency within the sequence. The histogram-based representation provides a compact and informative sequence summary, facilitating efficient computations and capturing sequence similarities.

The data used to construct a histogram are generated via a function *m*_*i*_ that counts the number of observations that fall into each disjoint category (bins). Thus, if we let *z* be the total number of observations and *k* be the total number of bins, the histogram data *m*_*i*_ meet the following conditions:

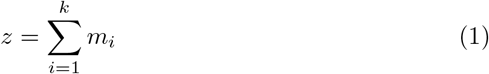

where *z* is the total number of *k*–mers in a sequence, and *m*_*i*_ represents the number of *k*–mers belonging to the *i*_*th*_ bin.

### 3.2 Gaussian Kernel Transformation

Once the histogram-based representations are obtained, Hist2Vec applies the Gaussian kernel transformation to map the histogram values into a high-dimensional feature space. The Gaussian kernel is a popular choice for capturing complex relationships between data points and is well-suited for capturing sequence similarities. The Gaussian kernel function is defined as:

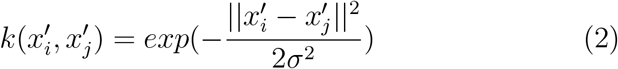

where 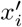 and 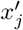 represent the histogram-based representations of sequence *x*_*i*_ and *x*_*j*_ respectively, and *σ* controls the width of the kernel. The kernel function measures the similarity between sequences in the high-dimensional feature space, with higher values indicating greater similarity.

Hist2Vec converts the histogram-based representations into a feature space where sequence similarities are kept by performing the Gaussian kernel transformation. This further helps apply standard machine learning (ML) algorithms to the modified feature space, leading to better classification results.

### 3.3 Kernel PCA for Embeddings

To further reduce the dimensionality and extract essential features, Hist2Vec employs kernel Principal Component Analysis (KPCA) to generate low-dimensional embeddings from the kernel matrix. Kernel PCA is a nonlinear extension of traditional PCA that operates in the feature space defined by the kernel function. The kernel matrix, denoted as *K*, is computed by applying the Gaussian kernel function to all pairs of histogram-based representations. Kernel PCA then performs eigendecomposition on the kernel matrix to extract the principal components, which capture the most crucial information. By selecting a subset of the principal components, Hist2Vec generates low-dimensional embeddings, which capture the similarities and variations between sequences in a compact representation, enabling efficient classification.

The algorithmic pseudocode for Hist2Vec is presented in Algorithm 1 and 2, while Figure 1 provides flowchart. The Algorithm 1 takes biological sequences and the total bins as input and calculates histogram embeddings based on the *k*-mers spectrum. First, the sequence is converted into *K*-mers (lines 2-5), then *K*-mers is converted into histogram embedding according to the number of bins (lines 6-9). While Algorithm 2 applies the Gaussian kernel to obtain the final kernel value (line 6). The kernel matrix is further processed for kernel-PCA and subsequently utilized in classification algorithms.

**Fig. 1:**
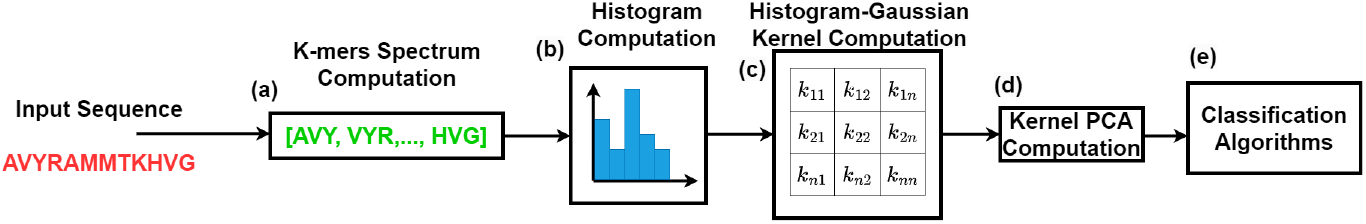
Workflow of Hist2Vec. The input represents a sequence vector computed using *k*-mers spectrum (a). The sequence is converted into the histogram (b) and kernel (c) matrix using His2Vec-Gaussian kernel, and then the kernel matrix is processed through Kernel-PCA (d) and used in the classification algorithms (e).

#### Algorithm 1 Histogram computation based on *k*-mers

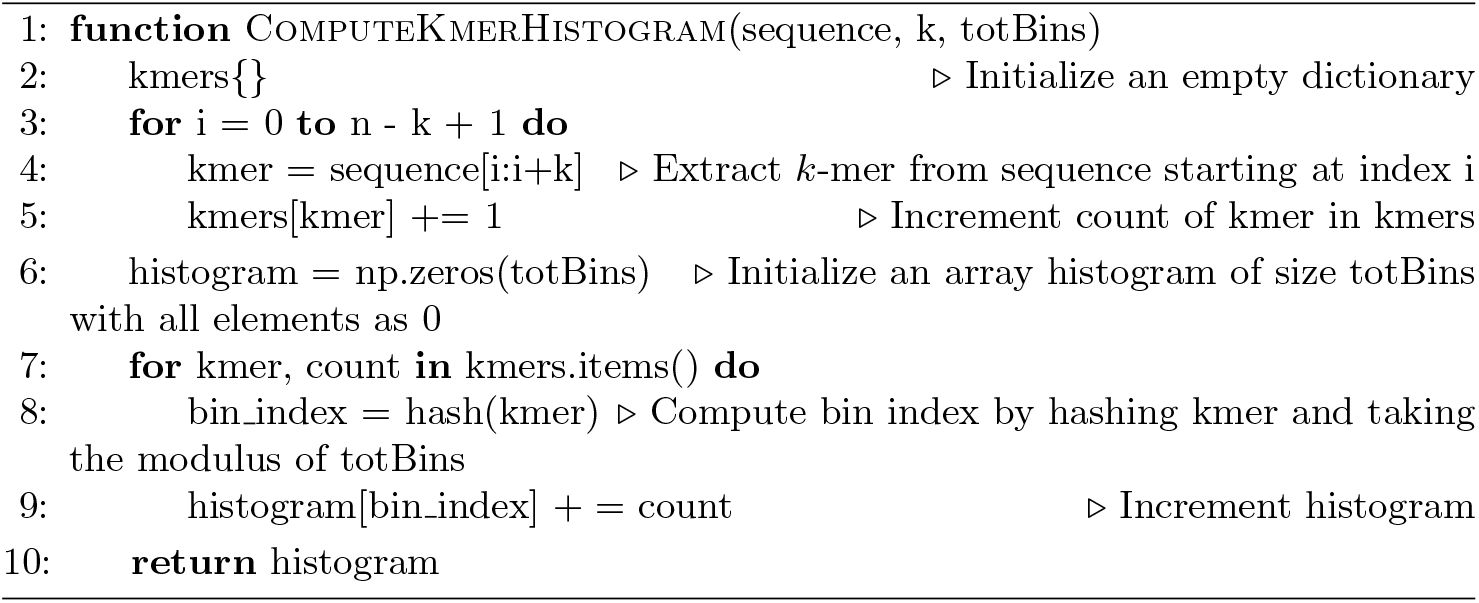

#### Algorithm 2 Hist2Vec-based kernel matrix generation

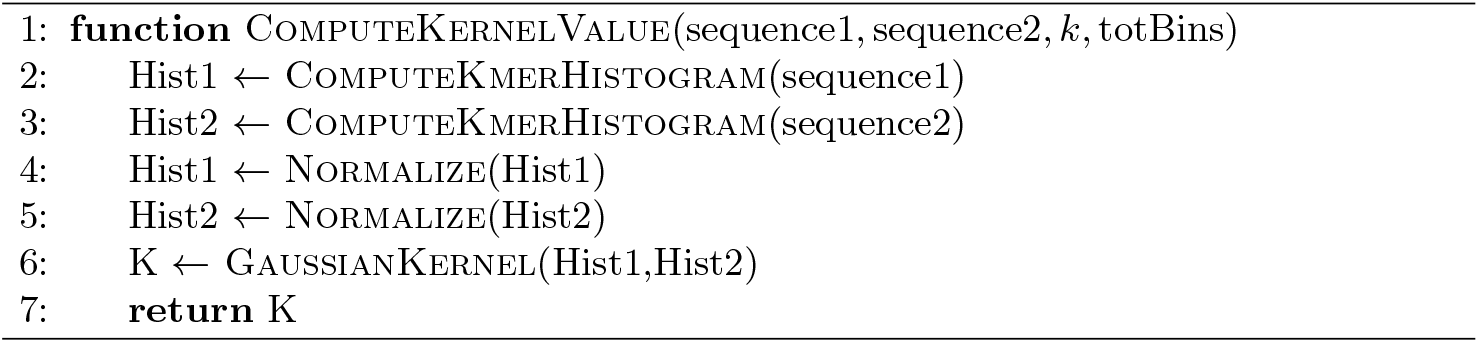

## 4 Experimental Setup

In this section, we detail the spike sequence dataset used for experimentation. We also discuss the baselines used for classification. In the end, we talk about the evaluation metrics used to test the performance of the models.

All experiments use an Intel(R) Core i5 system @ 2.10GHz having Windows 10 64 bit OS with 32 GB memory. For the classification algorithms, we use 70% of the data for training and 30% for testing. The 10% data from the training set is used as a validation set for hyperparameter tuning. We use two datasets to evaluate the performance of the proposed embedding method, which are explained below.

### Human DNA Dataset

This dataset uses the unaligned human DNA nucleotide sequences from [20]. It includes sequences and details on the relevant gene family for each sequence. It encodes data for seven distinct gene families (class labels). The count of sequences for each class label are G Protein Coupled (531), Tyrosine Kinase (534), Tyrosine Phosphatase (349), Synthetase (672), Synthase (711), Ion Channel (240), and Transcription Factor (1343) with the total count of 4380 sequences. The classification task classifies the gene family using the DNA sequence as input. Due to the variable length of sequences, since they are unaligned sequences, we have the maximum, minimum, and average lengths of 18921, 5, and 1263.59, respectively in our dataset.

### Coronavirus Host dataset

The spike sequences of the clades of the Coronaviridae family are extracted from ViPR [30,1] and GISAID ^3^, along with their metadata (genus/ subgenus, infected host, etc.), and the spike sequence and its corresponding host information are used to create our Coronavirus Host dataset. The number of hosts in our dataset is as follows: Bats (153), Bovines (88), Cats (123), Cattle (1), Equine (5), Fish (2), Humans (1813), Pangolins (21), Rats (26), Turtle (1), Weasel (994), Birds (374), Camels (297), Canis (40), Dolphins (7), Environment (1034), Hedgehog (15), Monkey (2), Python (2), Swines (558), and Unknown (2). We used 5558 spike sequences, which contain 21 unique hosts. Our classification jobs for this dataset use the hostname as the class label and sequences as input.

### Baseline Methods

To establish baselines, we choose newly suggested methods from several embedding generation categories, including feature engineering, conventional kernel matrix creation (including kernel PCA), neural networks, pretrained language models, and pre-trained transformers for protein sequences.

#### PWM2Vec

Feature Engineering method takes a biological sequence as input and designs fixed-length numerical embeddings [1].

#### String Kernel

Kernel Matrix-based method designs *n × n* kernel matrix that can be used with kernel classifiers or kernel PCA to get feature vector based on principal components [14,4].

#### WDGRL

A neural network (NN) based method takes the one-hot representation of biological sequence as input and designs an NN-based embedding method by minimizing loss [36].

#### AutoEncoder

This method uses a neural network (NN) to teach itself how to encode data as features. To iteratively optimize the objective, it applies the non-linear mapping approach. We used a 2 multilayer network with an ADAM optimizer and MSE loss function for our experiments [43].

#### SeqVec

It is a pre-trained Language Model which takes biological sequences as input and fine-tunes the weights based on a pre-trained model to get final embedding [18].

#### ProteinBert

It is a pre-trained Transformer, a protein sequence model to classify the given biological sequence using Transformer/Bert [9].

### 4.1 Evaluation Metrics and Classification Algorithms

We use average accuracy, precision, recall, F1 (weighted), F1 (macro), Receiver Operator Characteristic Curve (ROC), Area Under the Curve (AUC), and training runtime to compare the performance of various models. When using metrics created for the binary classification problem, we employ the one-vs-rest method for multi-class classification. Support Vector Machine (SVM), Naive Bayes (NB), Multi-Layer Perceptron (MLP), K-Nearest Neighbours (KNN), Random Forest (RF), Logistic Regression (LR), and Decision Tree (DT) are just a few of the linear and non-linear classifiers used in supervised analysis.

## 5 Results And Discussion

In Table 1, we present the classification results for the suggested and baseline methods for the **Human DNA** dataset. In terms of average accuracy, precision, recall, and F1 (weighted) and F1 (macro) scores, the Random Forest with Hist2Vec feature vectors outperformed all other baseline embedding approaches, while KNN using Hist2Vec-based embedding has the highest ROC-AUC score when compared to other embedding techniques. This suggests that the Hist2Vec embedding method is the method that performs the best for classifying sequences using ML models. Despite minimal train time, WDGRL’s classification performance is quite subpar. Compared to other embedding techniques, the WDGRL fails to preserve the overall data structure in its embeddings, which is one reason for this behavior. We can see that, except for one assessment criterion (training duration), Hist2Vec outperforms all baselines.

**Table 1:**
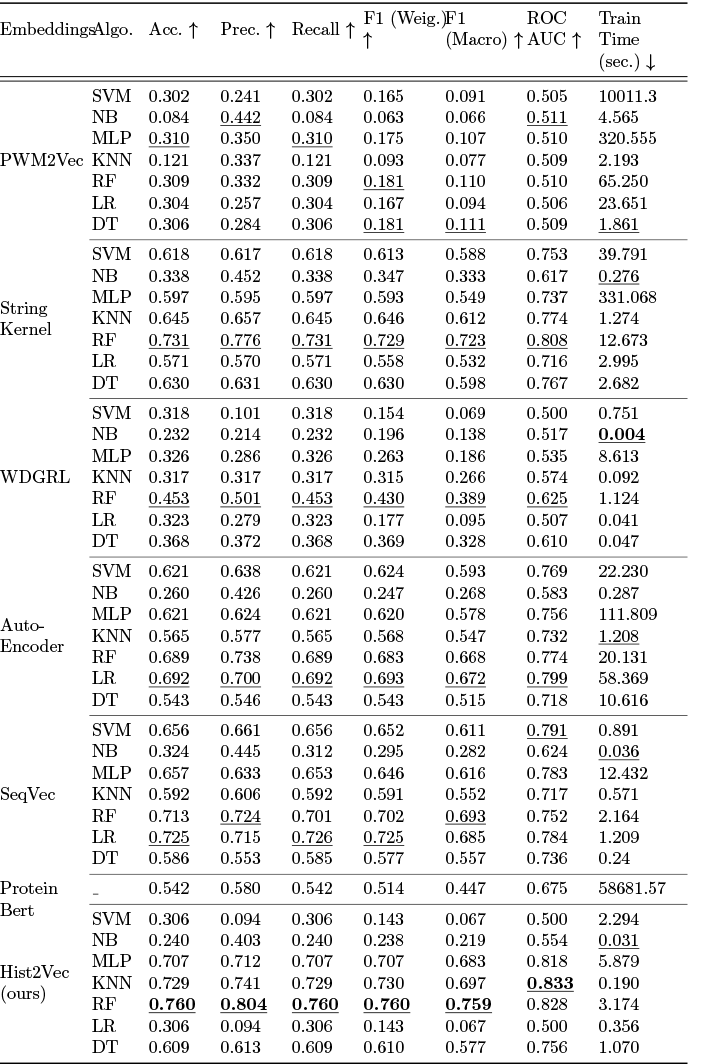
Classification results (averaged over 5 runs) on Human DNA dataset. The best classifier for respective embeddings is shown with the underline. Overall best values are shown in bold.

The results are presented in Table 2 for the **Coronavirus Host** dataset. Here also observe the performance of Hist2Vec on the host data sequences as compared to other baseline approaches. Because PWM2Vec is used to classify hosts (see [1]), we can compare Hist2Vec to PWM2Vec and other baselines to better understand its effectiveness. According to the results from the experiments, the RF classifier with Hist2Vec embedding works better than even PWM2Vec and different baselines in terms of accuracy, precision, recall, F1 weighted score, and AUC ROC score. Similarly, WDGRL has a low train time with the NB classifier in this instance, but the classifier performance is not even comparable. These findings suggest that the Hist2Vec method outperforms all other baseline techniques, including PWM2Vec.

**Table 2:**
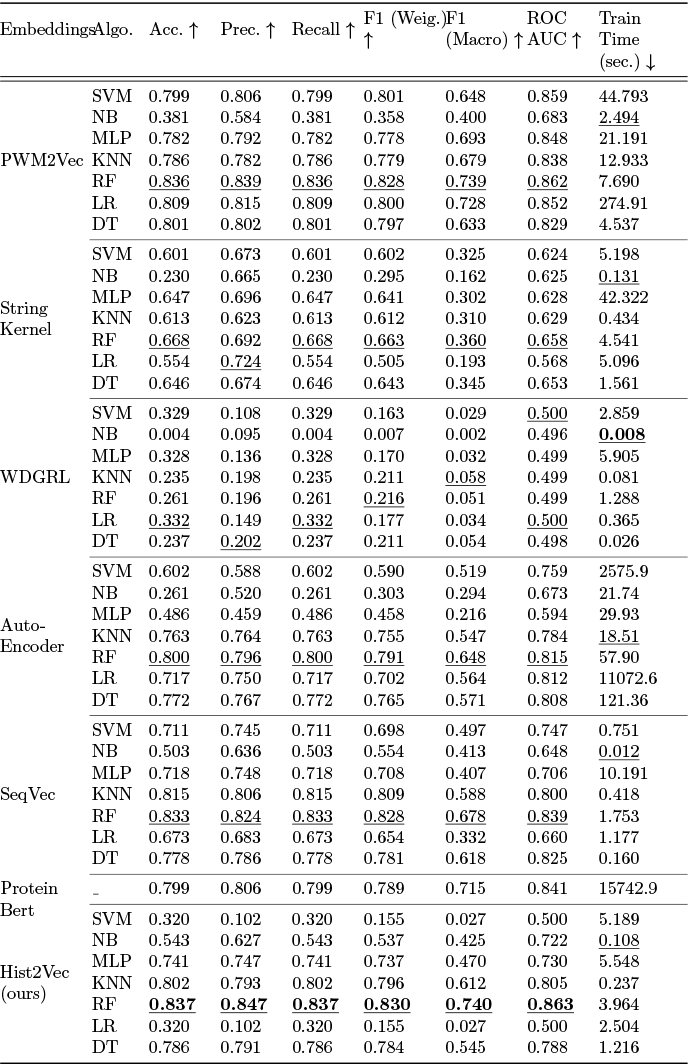
Classification results (averaged over 5 runs) on Coronavirus dataset. The best classifier for respective embeddings is shown with the underline. Overall best values are shown in bold.

## 6 Conclusion

We propose the Hist2Vec approach, an effective and alignment-free embedding method that outperforms state-of-the-art methods. Combining histogram-based embedding with a Gaussian kernel provides an efficient and effective framework for capturing sequence similarities and generating high-quality feature representations. Hist2Vec achieves the greatest accuracy of 76% and ROC AUC score of 83.3% for the classification of human DNA data and the highest accuracy of 83.7% and ROC AUC score of 86.3% for the classification of coronavirus hosts. Future studies involve assessing the Hist2Vec on other viruses, such as Zika.

## Footnotes

3 https://www.gisaid.org/

